# A genetically encoded ionic-stress sensor reveals protons as a sleep driver

**DOI:** 10.64898/2026.01.27.702131

**Authors:** Zhijian Ji, Junqiang Liu, Bingying Wang, Shuaian Wei, Yanchang Bian, Wenjie Zeng, Chan-I Chung, Zhimin Ma, Jianxiu Zhang, Xiaokun Shu, Dengke K. Ma

## Abstract

Dynamic ionic changes are hallmarks of physiological and behavioral state transitions, including sleep in animals. Although biosensors for specific cellular ions are widely available, real-time monitoring of overall ionic strength in living organisms remains challenging. Here, we present a <u>g</u>enetically <u>e</u>ncoded <u>n</u>uclear translocation ionic <u>s</u>ensor (GENTIS) that enables direct visualization of ionic stress *in vivo*. Using GENTIS in *C. elegans*, we uncover rhythmic elevations in ionic strength during larval molting transitions that coincide with the lethargus sleep. Cytosolic proton ionic increase through inhibition of v-ATPase is sufficient to induce GENTIS nuclear translocation and evoke behavioral quiescence, characterized by reduced feeding and activation of sleep-active neurons. Apical membrane v-ATPases undergo disassembly during lethargus and under sleep-inducing stress conditions, leading to proton accumulation. Notably, this proton-mediated sleep is suppressed by proton buffering with ammonium. Together, these findings establish GENTIS as a powerful tool for tracking ionic strength dynamics in living organisms and support protons as a physiological driver of sleep.

## Introduction

Sleep is an evolutionarily conserved behavioral state essential for animal survival and physiological homeostasis(*1–7*). In *Caenorhabditis elegans*, larvae enter a quiescent sleep state during each molting cycle, referred to as lethargus(*8–10*). Lethargus is characterized by cessation of feeding and locomotion and displays the hallmarks of sleep: reduced responsiveness, reversibility, and homeostatic regulation(*10–13*). The GABAergic and peptidergic RIS neuron has been identified as a key sleep-active neuron in this developmental sleep, becoming strongly activated at sleep onset and driving behavioral quiescence through neuropeptide signaling(*14*). In adult *C. elegans*, various types of stress can also induce sleep mediated primarily by the ALA neuron, which responds to cellular damage signals by releasing FMRFamide-like neuropeptides to induce a reversible sleep-like state(*15–17*). RIS and ALA neurons can also act coordinately in a neural circuit driving sleep behaviors, including pumping and locomotion quiescence(*18*). Neuropeptides released by ALA and RIS act on the receptor DMSR-1 that induces and self-inhibits sleep for limiting the duration of sleep(*19*, *20*). Both developmental and stress-induced sleep appear to involve EGF signaling: overexpression of the EGF ligand such as LIN-3 and SISS-1 can promote quiescence, and stress-induced sleep requires EGF pathway components(*16*, *21*, *22*). However, the physiological signals that link developmental cues, metabolic state or cellular stress to sleep still remain incompletely understood.

Emerging studies in mammals and invertebrates suggest that sleep is intimately tied to metabolic and ionic homeostasis(*23–25*). Changes in the local concentration of protons, the smallest and simplest ionic species, can profoundly influence cellular physiology. Proton gradients are established across cellular membranes, including those of mitochondria, lysosomes, and epithelial apical domains, through the coordinated activity of membrane-bound proton pumps, exchangers and transporters that couple chemical energy, such as ATP hydrolysis or redox reactions, to directional proton translocation(*26–28*). In *Drosophila*, reduced energy demand in sleep-active neurons leads to the accumulation of protons and electrons, forming a biochemical basis for sleep pressure(*29*). Moreover, intracellular acidification and local proton flux have been implicated in modulating neuronal excitability and synaptic plasticity in vertebrates, indicating that protons may serve as conserved physiological messengers coupling metabolic state to neural circuit activity(*30*). In *C. elegans*, gut-derived protons function as paracrine transmitter signals that drive muscle contraction during the rhythmic defecation cycle(*31*). However, whether proton dynamics occur during sleep transitions and contribute to signaling to sleep-regulatory circuits remains unknown.

To monitor ionic strength *in vivo*, existing genetically encoded ion sensors, such as pHluorin for protons and other fluorescence-based probes, are widely useful yet limited either in their intrinsically limited dynamic range and modest signal amplitude, or can typically detect specific ions or large changes within restricted subcellular compartments(*32–34*). To overcome these limitations, we developed a genetically encoded nuclear translocation ionic sensor. This sensor (named as GENTIS, see below) is an engineered fusion protein whose nuclear import, rather than intrinsic fluorescence change, is sensitively activated by elevated ionic strength. Using GENTIS, we found that *C. elegans* larvae experience rhythmic ionic stress during each larval molt, coinciding with periods of feeding quiescence. Moreover, pharmacological inhibition of proton-pumping v-ATPases induces ionic stress and activates sleep behavior via the known sleep-promoting neurons and an intestinal H⁺/Na⁺ exchanger. Together, our findings identify protons as a physiological driver of sleep and establish GENTIS as a powerful tool for visualizing ionic stress dynamics in living organisms.

## Results

### Design and validation of GENTIS

We rationally engineered a <u>g</u>enetically <u>e</u>ncoded <u>n</u>uclear translocation ionic <u>s</u>ensor (GENTIS) by inserting two oppositely charged α-helical domains, whose intrahelical salt bridges respond to ionic strength(*35*), between a classical nuclear localization signal (NLS) and a nuclear export signal (NES) (Fig. 1A). Under low-salt conditions, electrostatic interactions within the helix stabilize a conformation that masks the NLS and favors cytosolic localization through the opposing NES. When ionic strength increases, these salt bridges are weakened, exposing the NLS and promoting importin-dependent nuclear import. After optimizing construct design and concentration for germline micro-injection, we generated stable transgenic *C. elegans* strains expressing cytosolic GENTIS under the control of ubiquitously active *rpl-28* promoter. Exposure to known ionic stress-increasing conditions, including hypertonic media such as 10% sorbitol, 10% glucose, or high extracellular ionic conditions (250 mM NaCl or 250 mM NH_4_OAc), triggered a striking relocalization of the fluorescent reporter of GENTIS from the cytosol to the nucleus (Fig. 1B). The GENTIS translocation response was dose-dependent and peaked within the first 20 minutes (Fig. 1C), indicating that GENTIS sensitively reports ionic strength changes with high temporal resolution. We observed GENTIS translocation upon exogenous hypertonic or ionic stresses most strikingly in the intestine and consistently across all larval stages (Fig. S1A).

**Fig. 1.**
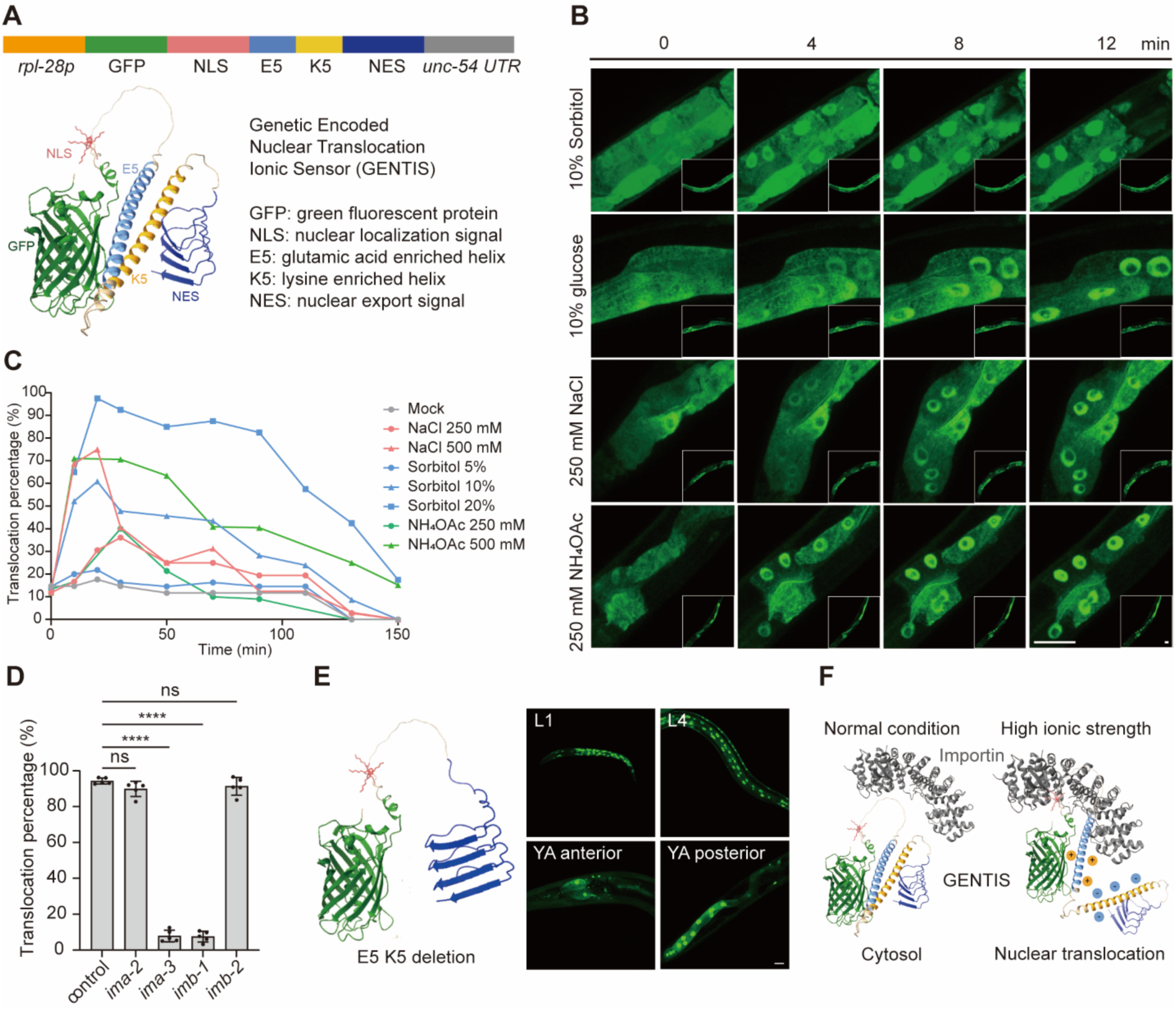
Development of a genetically encoded nuclear translocation ionic sensor. (A) Schematic diagram illustrating the design of the GENTIS sensor. (B) Representative fluorescence images of animals showing GENTIS translocation from the cytosol to the nucleus in response to hypertonic or high ionic strength conditions. (C) Dose-dependent response and time-course of GENTIS nuclear translocation under the indicated stress conditions. (D) GENTIS translocation requires the importin-dependent nuclear import machinery. **** indicates P<0.001 (n = 5 independent trials). (E) Representative images of worms expressing a GENTIS variant lacking the oppositely charged helix (E5 and K5), showing constitutive nuclear localization of the sensor without ionic stress sensitivity. (F) Proposed model of GENTIS translocation mechanism and behavior under high ionic strength conditions. Scale bars, 20 μm.

We next examined GENTIS responses under importin-inhibited states and additional stress conditions. Disruption of the classical nuclear import machinery, for example through RNAi against genes encoding key importin machinery, abolished GENTIS translocation under high ionic stress, confirming its dependence on the importin pathway (Fig. 1D). A control reporter lacking the charged helix showed constitutive nuclear localization even under unstressed conditions, supporting the helix’s role in masking the NLS (Fig. 1E). GENTIS underwent nuclear translocation after UV irradiation and acute heat stress but not oxidative stress, and the kinetics were slower than those observed under ionic stress (Fig. S1B), suggesting distinct underlying mechanisms. GENTIS also responded similarly to different metal ion treatments, showing no preference for specific ions (Fig. S1C, 1D), indicating that it broadly senses hypertonicity-induced or direct ionic stress types rather than distinct ion species.

Together, these results establish GENTIS as a robust, genetically encoded reporter of intracellular ionic strength that readouts facile translocation between the cytosol and nucleus through balanced NLS–NES regulation in response to ionic stress (Fig. 1F).

### GENTIS translocation is independent of protein phase separation

Hyperosmotic stress can induce phase separation or aggregation of proteins in other systems(*36*). To test whether GENTIS movement reflects cellular phase separation, we generated a transgenic strain expressing a known phase-separation reporter (SPAR) and applied a commonly used liquid–liquid phase separation (LLPS) inhibitor 1,6-hexanediol(*37*, *38*). As expected, hypertonic stress induced visible SPAR phase-separated condensates, which were abolished by 1,6-hexanediol (Fig. S2A). However, we observed that GENTIS still translocated to the nucleus under hypertonic conditions in the presence of 1,6-hexanediol (Fig. S2B). Thus, liquid-liquid phase separation is unlikely to cause GENTIS nuclear translocation; rather, GENTIS directly senses ionic strength changes via the designed helices carrying opposite charges.

### Developmental ionic stress coincides with molting sleep

We next examined endogenous GENTIS dynamics during natural larval development. During the L2 and L3 stages, we observed transient nuclear localization of GENTIS in many animals, with marked peaks around the expected times of lethargus (Fig. 2A–C). Quantification of populations of animals confirmed that the fraction showing nuclear GENTIS oscillated in phase with the lethargus periods during molting (Fig. 2B). Time-lapse imaging of individual animals revealed that GENTIS translocation occurred rhythmically, peaking at each larval molt lethargus (Fig. 2C, 2D). These observations indicate a recurring rise in internal ionic strength during the molts. To correlate GENTIS with sleep behavior, we synchronized animals and simultaneously monitored pharyngeal pumping. GENTIS nuclear translocation onset occurred at the L1–L2 transition, concomitant with a sharp decline in pumping (Fig. 2F). Across animals, those with more nuclei showing GENTIS translocation exhibited markedly lower pumping rates (Fig. 2G). In a bead-feeding assay, animals displaying GENTIS nuclear localization ingested far fewer beads than non-translocated controls (Fig. 2H), confirming reduced feeding during the lethargus sleep. We found that GENTIS still translocated into nuclei during the L1–L2 transition when animals were incubated in external hypotonic solution (pure water) (Fig. S3A). In addition, the intestinal nuclear envelope and cellular organelles including lysosomes remained largely intact during the L1–L2 transition, as indicated by various specific markers (Fig. S3B, 3C). Together, these data reveal that *C. elegans* larvae undergo pulses of ionic stress in each molting cycle, which coincide with behavioral quiescence during the lethargus sleep at molts.

**Fig. 2.**
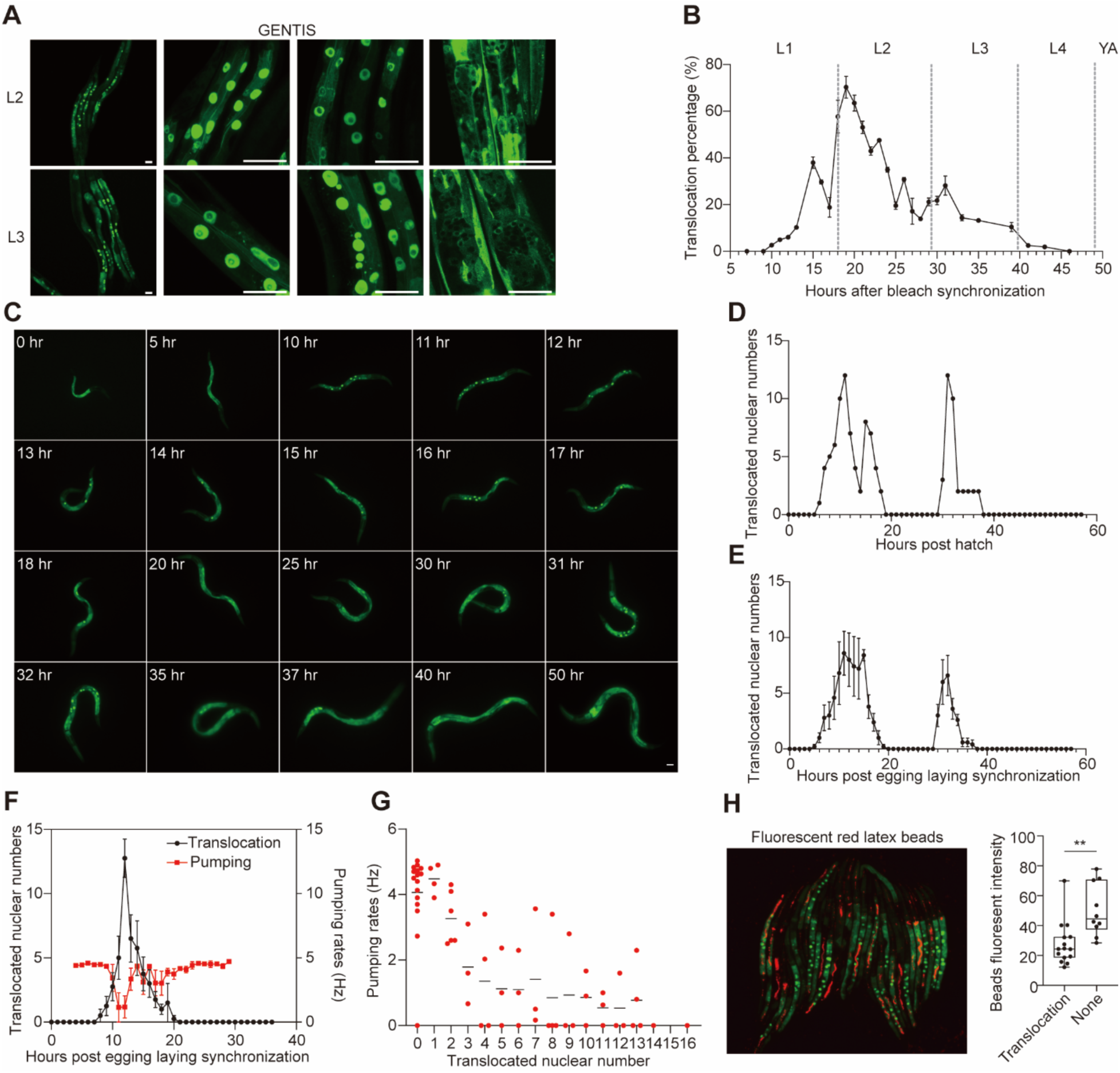
GENTIS reveals a natural ionic stress response during larval development. (A) Representative confocal images of L2- and L3-stage larvae expressing GENTIS, showing nuclear translocation of the sensor at these developmental stages. (B) Percentage of worms showing GENTIS nuclear translocation during larval development. n > 30 worms were analyzed per condition, and experiments were repeated independently three times. Note the population is not strictly synchronized by the bleach protocol. (C) Time-lapse fluorescence images showing rhythmic GENTIS nuclear translocation during early larval development (n=3). (D) Time-course of the number of nuclei with GENTIS translocation in the individual animal shown in (C), illustrating the dynamics of nuclear translocation during development. (E) Nuclear translocation kinetics during larval development for animals synchronized by egg-laying (n = 5). (F) Simultaneous tracking of pharyngeal pumping and GENTIS nuclear translocation in animals during the L1–L2 larval transition (n=4). (G) Quantification of pharyngeal pumping rates in animals with different numbers of GENTIS-translocated nuclei. Scale bars, 20 μm. (H) Representative images and quantification of worms fed with red fluorescent latex beads for 10 min, comparing feeding (pharyngeal intake) between animals with and without GENTIS translocation. ** indicates P<0.01 (n >10 independent animals).

### v-ATPase inhibition is sufficient to induce proton stress and sleep

To identify molecular components required for GENTIS activation under ionic stress, we performed targeted RNAi and pharmacological inhibition of candidate ion transport and signaling pathways (Fig. S4A, 4B). We systematically knocked down representatives of major channel and transporter families, including aquaporins (*aqp-1*, *aqp-2*, *aqp-4*, *aqp-8*), Na⁺/K⁺-ATPases and cotransporters (*eat-6*, *nkcc-1*), chloride and cation channels (*clh-6*, *cup-5*), and regulators of ion homeostasis (*wnk-1*, *cul-3)* highly expressed in the intestine. Although RNAi efficiency for these genes varies and remains fully validated, none of these perturbations in single RNAi treatment prevented GENTIS nuclear translocation in response to hypertonic stress. By contrast, RNAi against genes encoding importin machineries strongly attenuated GENTIS translocation (Fig. 1D).

Pharmacological inhibition of ion-regulatory signaling using the pan-WNK kinase inhibitor WNK463 also did not alter GENTIS dynamics. Although GENTIS showed the strongest translocation in CuSO_4_ among all metal ions tested (Fig. S1C), depletion of copper transport and homeostasis genes (*cua-1*, *cuc-1*, *chca-1*, *F21D5.3*) had little effect. Likewise, RNAi of TOR pathway regulators (*let-363*, *daf-15*) was ineffective.

To complement the RNAi analysis, we systematically screened a panel of small molecules targeting ion transport and proton regulatory pathways (Fig. S4B). Inhibitors of Na⁺, K⁺, and Cl⁻ channels, including amiloride (Na⁺ channel blocker), glibenclamide (ATP-sensitive K⁺ channel blocker), bumetanide (NKCC cotransporter inhibitor), NPPB and DIDS (chloride channel and anion exchange inhibitors), as well as a chloride ionophore cocktail, had no effect on GENTIS nuclear translocation. Similarly, calcium channel blockers and ion regulators such as ruthenium red and lanthanum chloride, and the intracellular calcium chelator BAPTA-AM, failed to suppress GENTIS reporter translocation. Chelation with EDTA also had no inhibitory effect, indicating that general sequestration of divalent cations does not modulate the sensor.

We next examined proton pump inhibitors. Diphyllin and omeprazole, which target v-ATPase and H⁺/K⁺-ATPase activity in mammalian cells, respectively, did not affect GENTIS translocation under baseline or hypertonic stress conditions. In contrast, we found that 4-(2-Furyl)-3-buten-2-one (furfural acetone, FA), a previously reported *C. elegans* v-ATPase (*vha-12* and *vha-13*) inhibitor(*39*), robustly induced nuclear translocation of GENTIS in a dose-dependent manner, mimicking the ionic stress response and likely resulting from elevated cytosolic proton levels (Fig. 3A, 3B).

**Fig. 3.**
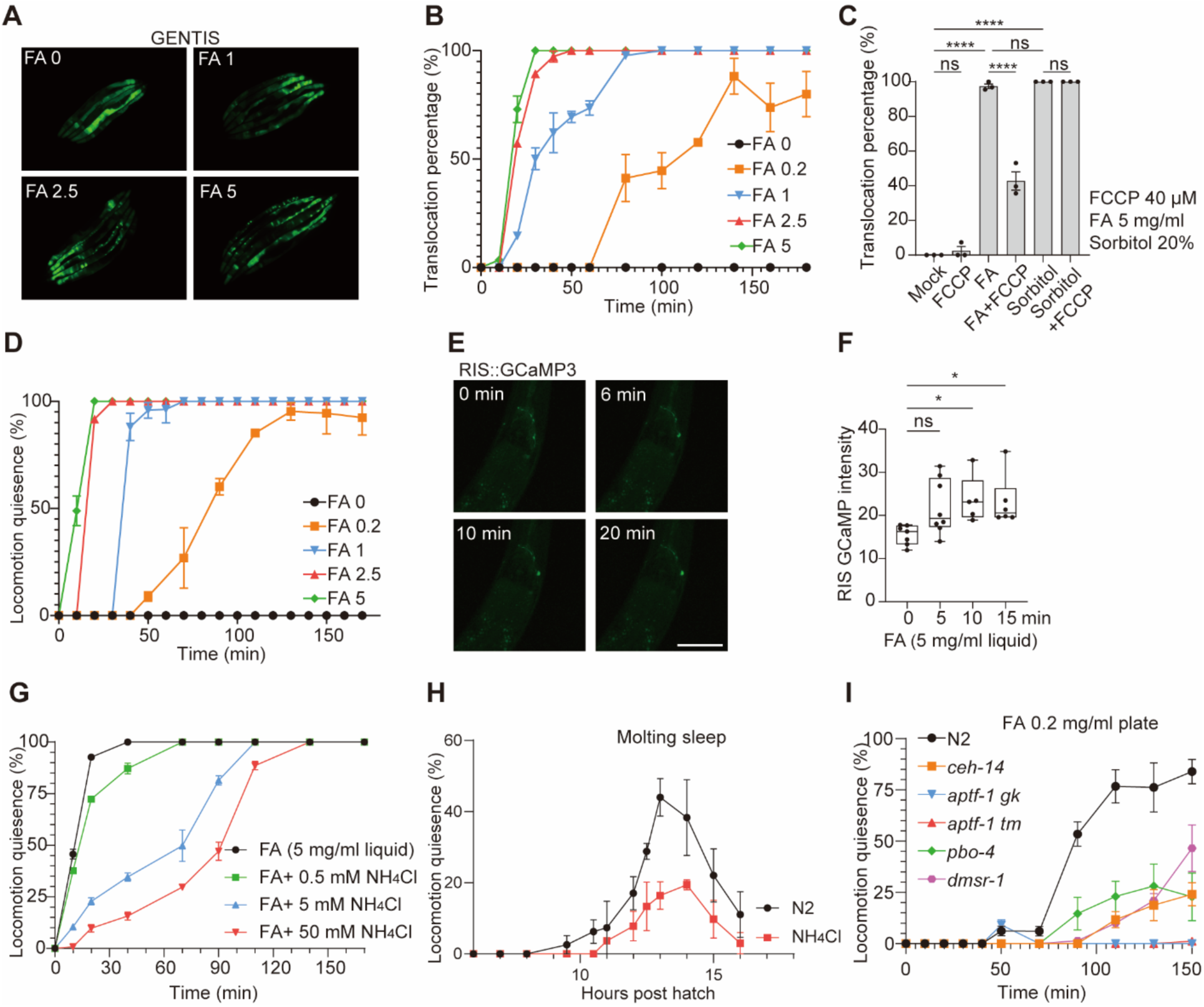
Coupling of proton-driven ionic stress to sleep induction by VHA inhibitor. (A) Representative fluorescence images showing GENTIS nuclear translocation in young adult animals treated with FA in plates at the indicated doses. (B) Time course of GENTIS translocation percentage in response to increasing FA doses in plates. (C) GENTIS translocation is suppressed by the protonophore FCCP (40 μM) when co-treated with FA (5 mg/ml in M9) for 10 min, but not when co-treated with sorbitol (20%), indicating a proton-specific effect of FA. **** indicates P<0.001 (n = 3 independent trials). (D) Fraction of worms entering sleep state (locomotion quiescence) after FA treatment at the indicated doses in plates. (E and F) Activation of the sleep-promoting RIS neuron by FA (5 mg/ml in M9), as indicated by increased calcium-dependent fluorescence in RIS::GCaMP animals over time. * indicates P<0.05 (n > 5 independent animals). (G) Sleep induction by FA (5 mg/ml in M9) is dose-dependently suppressed by co-treatment with NH_4_Cl, a membrane-permeable weak base. (H) Fraction of animals exhibiting locomotion quiescence during the L1–L2 transition under NH_4_Cl or mock treatment. (I) FA-induced pharyngeal pumping quiescence (sleep) is dependent on the ALA and RIS neurons and a H^+^/Na^+^ exchanger (*pbo-4*). n > 20 animals were analyzed per condition, and experiments were repeated independently three times. Error bars represent S.E.M.

To determine whether cytosolic proton accumulation upon FA treatment contributes to GENTIS activation, we co-treated animals with the protonophore FCCP, which reduces cytosolic proton concentration by facilitating proton flux into mitochondria. 40 μM FCCP treatment markedly suppressed FA–induced GENTIS nuclear translocation but did not affect sorbitol-induced translocation (Fig. 3C). These results indicate that FA triggers ionic stress and GENTIS translocation primarily through v-ATPase inhibition and cytosolic proton elevation, whereas sorbitol acts via osmotic-ionic mechanisms independent of proton accumulation.

Remarkably, the same doses that triggered GENTIS translocation also caused behavioral locomotion quiescence, with kinetics similar to natural sleep bouts (Fig. 3D). The FA-induced quiescence was reversible: animals resumed pumping rapidly after brief exposure (Fig. S5A), indicating a *bona fide* sleep state. Concurrent imaging of the sleep-active RIS neuron (GCaMP in RIS) showed robust Ca^2+^ activation of RIS following FA (Fig. 3E, 3F), consistent with engagement of sleep-promoting neurons, rather than non-specific paralytic effects on pharyngeal or body wall muscles.

### Proton accumulation is required for sleep induction

To further test the role of protons in FA-induced sleep, we co-treated animals with ammonium chloride (NH_4_Cl), a membrane-permeable weak base that buffers intracellular protons. NH_4_Cl dose-dependently suppressed FA-induced sleep but not sorbitol induced sleep (Fig. 3G, S5B, S5D), with high doses almost completely blocking quiescence despite continued FA exposure. In contrast, GENTIS nuclear translocation remained unaffected by NH_4_Cl co-treatment (Fig. S5C), indicating that NH_4_Cl does not prevent FA-induced ionic stress *per se*, but rather counteracts proton accumulation. To examine how proton buffering influences sleep dynamics during larval development, we tracked animals treated with NH_4_Cl through the L1–L2 transition. NH_4_Cl had little effect on the onset of developmental pumping quiescence during L1-L2 lethargus (Fig. S5F), but it markedly reduced the fraction of animals entering locomotion quiescence (Fig. 3H), indicating that proton accumulation contributes specifically to the coordination of full behavioral sleep. We also quantified the ratio of locomotion-to-pumping quiescence to determine the proportion of animals exhibiting locomotion sleep among those that entered pumping quiescence, thereby controlling for stage-dependent or other factors that may prevent some animals from sleeping. Consistently, we found that NH_4_Cl treatment also decreased the locomotion-to-pumping quiescence ratio (Fig. S5K). Together, these results indicate that intracellular proton accumulation is a critical determinant of FA-induced and developmental sleep.

To define the neural and molecular components mediating FA-induced and developmental sleep, we examined mutants defective in the ALA and RIS neurons as well as in proton and neuropeptide signaling pathways. Wild-type animals displayed robust sleep-related behavioral quiescence after FA treatment (0.2 mg/ml), whereas *aptf-1* mutants lacking the sleep-active RIS neuron completely failed to sleep (Fig. 3I). *ceh-14*, *dmsr-1*, and *pbo-4* mutants entered only partial quiescence, indicating that ALA neuron, the *dmsr-1* receptor, and intestinal H^+^/Na^+^ exchange via *pbo-4* are required for full FA-induced sleep (Fig. 3I). By contrast, *pbo-5*, *adm-4*, and *siss-1* mutants showed sleep comparable to wild-type animals, suggesting that cholinergic signaling, metalloprotease, and EGF-like ligand release are not essential for this process (Fig. S5E). During developmental lethargus, while wild-type animals showed tightly coupled suppression of both locomotion and pumping, *aptf-1*, *ceh-14*, *dmsr-1*, and *pbo-4* mutants exhibited reduced locomotion quiescence during the L1–L2 molt (Fig. S5G). This was accompanied by delayed or incomplete suppression of pharyngeal pumping (Fig. S5H, S5I). *lin-42* mutants, which disrupt the molting timer, exhibited unusually longer sleep as measured by pumping (Fig. S5H). Quantification of the locomotion-to-pumping quiescence ratio confirmed that wild-type animals exhibit tightly coupled behavioral quiescence, whereas *aptf-1* mutants display a nearly complete loss of locomotion quiescence despite suppression of pumping (Fig. S5J).

### Regulation of v-ATPase during developmental and stress-induced sleep

We further investigated the effect of FA on intestinal proton transport. Although FA has been reported to inhibit the v-ATPase subunits VHA-12 and VHA-13, structural modeling suggests that FA may bind near the ATP-binding site of VHA-13 (Fig. 4A). We monitored the effect of FA exposure on VHA-13 endogenously tagged with the split mScarlet-complemented fluorescent protein by CRISPR knock-in(*40*). FA treatment caused a marked loss of the intestinal apical mScarlet::VHA-13 fluorescence over time (Fig. 4B, 4C). Surprisingly, mScarlet::VHA-13 signal also declined under other stress conditions, including UV irradiation (0.1 J/m²) and 20% sorbitol treatment (Fig. S5L). The intestinal apical membrane remained intact under all these conditions, as indicated by unaltered OPT-2::GFP signal (Fig. S5L). Notably, endogenous mScarlet::VHA-13 levels at the apical surface also decreased during developmental lethargus (15-16.5 hrs post hatch) in larvae (Fig. 4D-4F), suggesting that v-ATPase activity is normally downregulated during molting sleep. Moreover, we observed that FA-induced sleep was reduced in *vha-13* gain-of-function (GOF) mutants compared to wild-type animals (Fig. S5M). Together, our findings demonstrate that pharmacological inhibition of v-ATPase by FA recapitulates a physiological feature of sleep, and the consistent and robust VHA-13 reduction across diverse sleep-inducing stress conditions suggests a widespread mechanism whereby organisms can downregulate v-ATPase activity and increase proton flux to drive sleep behaviors (Fig. 4G).

**Fig. 4.**
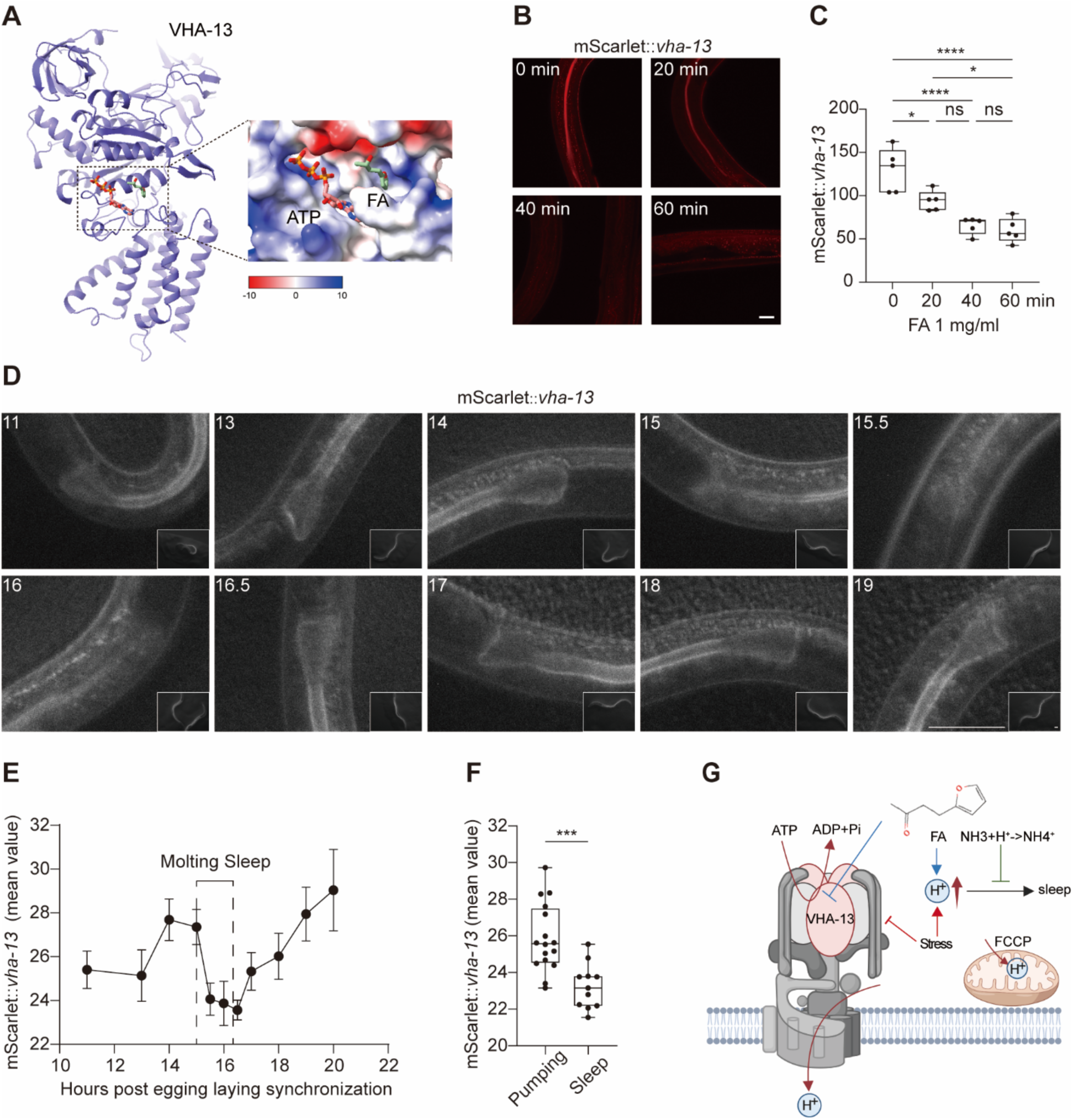
Downregulation of VHA-13 links intestinal proton transport to sleep induction. (A) Structural modeling of FA binding to the ATP-binding pocket of the VHA-13 subunit. Structural analysis predicts that FA occupies a site adjacent to the ATP-binding region, potentially impacting nucleotide binding and disrupting proton translocation (red, −10 kcal·mol⁻¹·e⁻¹; white, 0; blue, +10 kcal·mol⁻¹·e⁻¹). (B) Representative images showing redistribution of intestinal mScarlet::VHA-13 following 1 mg/ ml FA treatment. Apical membrane signal is progressively reduced over time. Scale bar, 20 μm. (C) Quantification of apical mScarlet::VHA-13 fluorescence intensity over a 60-minute time course after FA exposure (1 mg/ml). **** indicates P<0.001, * indicates P<0.05 (n > 5 independent trials). (D) Time-lapse imaging of anterior intestinal mScarlet::VHA-13 expression during the L1–L2 molt. Time points (in hours post-hatching) are indicated; lethargus occurs between 15 and 16.5 hours. Insets show full-field views before zoom-in. Scale bars, 20 μm. (E) Quantification of mScarlet::VHA-13 signal during development, showing a transient decline in apical fluorescence during molting sleep (n = 5 animals). (F) Comparison of mScarlet::VHA-13 intensity between animals in active feeding (pumping) and sleep states, revealing significantly lower apical levels during sleep. *** indicates P<0.005 (n > 10 independent trials). (G) Schematic model of proton-mediated sleep regulation. Inhibition of VHA-13 by FA and other stresses leads to reduced proton export and cytosolic proton accumulation, triggering sleep. FCCP redistributes protons into mitochondria suppressing GENTIS translocation, and NH_4_Cl buffers intracellular protons suppressing FA-induced sleep.

## Discussion

Our results identify intracellular proton accumulation as a physiological signal coupling metabolic state and ionic stress to sleep. During both developmental and stress-induced sleep, inhibition of proton pumping by intestinal VHA-13 coincides with and is causally linked to elevated cytosolic proton concentration and feeding quiescence. This link between proton buildup and sleep is further supported by recent findings in *Drosophila*, where photo-energized mitochondrial proton/H⁺ outflux into the cytosol of sleep-active dFB neurons increases sleep drive(*29*). Importantly, discharging proton motive force by the proton leak channel Ucp4 attenuates sleep drive in *Drosophila* (*29*). Elevated proton accumulation in the mitochondrial intermembrane space or cytosol can inhibit respiratory electron transport, increasing reactive oxygen species (ROS) that may signal or promote sleep(*41*). As metabolic buildup and sleep deprivation can increase tissue acidity and subsequent ROS generation, moderate ROS may act as physiological signals that promote sleep, whereas excessive ROS can trigger inflammation and tissue damage(*42–44*). In *C. elegans*, intestinal apical v-ATPase activity supports nutrient uptake by powering proton-coupled transporters; its downregulation during lethargus likely reflects a coordinated suppression of digestion and anabolic metabolism linked to behavioral quiescence(*45–47*). Our genetic and pharmacological manipulations demonstrate that v-ATPase activity slowdown and sleep are not merely correlated but mechanistically intertwined through the regulation of intracellular proton flux.

Across animals, proton gradients are central to bioenergetic homeostasis. In yeast, nutrient deprivation triggers v-ATPase disassembly from vacuolar membranes, diminishing proton transport in response to glucose limitation(*48*, *49*). Shifts in proton concentration accompany metabolic suppression during mammalian torpor and hibernation, where tissue acidification and reduced transmembrane proton flux are hallmarks of energy conservation(*50*). The parallel downregulation of proton pumping thus mirrors broader cross-kingdom responses to energetic stress. By enabling real-time visualization of ionic stress *in vivo*, GENTIS reveals that proton accumulation can serve as a potent cue coupling cell bioenergetic status to the initiation of *C. elegans* sleep quiescence. More broadly, these findings define a conceptual framework in which protons serve as a conserved biochemical switch that synchronizes cellular energy and ionic balance with organismal quiescence, from sleep to other dormancy states.

## Methods

### Sensor construction

The GENTIS coding sequence was assembled by inserting two designed helices with acidic (E5 helix-5X EVSALEK) and basic residues (K5 helix-5X KVSALKE) separately between a classical NLS (PKKKRKV) and 4XNES (PLQLPPLERLTLSQDGGSAGG). A GFP fluorescent protein in the N terminal was included to visualize localization. A control variant lacking the charged helix was constructed similarly. All PCR constructs were expressed in *C. elegans* under the ubiquitous promoter *rpl-28*p and *unc-54*utr via microinjection.

### *C. elegans* culture and imaging

*C. elegans* strains were maintained using standard protocols unless other specified. The N2 Bristol strain was used as the reference wild type(*51*). The GENTIS construct was micro-injected to the *C. elegans* germline to form extrachromosomal arrays followed by UV-mediated integration and the strain was N2 back-crossed at least three times. Genotypes of mutant or transgenic strains used are as follows: VC1669 *aptf-1(gk794)* II, CVB96 *siss-1(csn20)* IV, MLC1778 *vha-13(luc133gof)* V, OH15422 *ceh-14(ot900)* X, CZ2744 *adm-4(ok265)* X, MT6929 *pbo-4(n2658)* X, DMS2851 *dmaIs185 [rpl-28p::GENTIS; unc-54p::mCherry]* II, DMS852 *dmaIs30 [SPAR; unc-54p::mCherry],* HBR2727 *dmsr-1(qn45)* V; MT5936 *pbo-5(n2303)* V*, goeIs304[flp-11p::SL1-GCaMP3.35-SL2::mKate2-unc-54UTR, unc-119(+)],* HBR1361 *goeIs304[flp-11p::SL1-GCaMP3.35-SL2::mKate2-unc-54UTR, unc-119(+)],* CF4586 *muIs252 II; unc-119(ed3)* III*; vha-13(mu493[wrmScarlet11::vha-13])* V.

For GENTIS population assays, synchronized animals were transferred to NGM plates and exposed to solutions containing sorbitol, NaCl, or other salts at the indicated concentrations. For individual pharyngeal pumping analysis, synchronized L1 larvae were placed on thin-layer NGM plates and monitored using an EVOS M5000 microscope. Confocal and compound fluorescence images were acquired using Leica systems to assess GENTIS localization. For time-course experiments, live GENTIS-expressing animals were mounted on agarose pads and imaged at defined intervals following solution treatment. The CRISPR knock-in mScarlet::VHA-13 animals were cultured on thin-layer NGM plates and imaged under the EVOS M5000 system. For FA experiments, animals were treated with 4-(2-Furyl)-3-buten-2-one (FA) (TCI, F0235) solution poured on the plates or the plates containing FA.

SPE confocal (Leica) and digital automated epifluorescence microscopes (EVOS, Life Technologies) were used to capture fluorescence images. Animals were randomly picked at the same stage and treated with 1 mM levamisole in M9 solution, aligned on a 2% agar pad on a slide for imaging. Identical setting and conditions were used to compare experimental groups with control. For quantification of GFP fluorescence, animals were outlined and quantified by measuring gray values using the ImageJ software. The data were plotted and analyzed by using GraphPad Prism10.

### Phase separation tests

Transgenic animals expressing a known phase separation and condensation-based reporter (SPAR)(*38*) were exposed to hypertonic shock (e.g. 500 mM NaCl or 20% sorbitol) with or without 5% or 10% 1,6-hexanediol. Condensate formation and GENTIS localization were imaged by confocal microscopy.

### Behavioral assays

Molting-related quiescence was monitored in L1 larvae from egg-laying synchronization by tracking pharyngeal pumping in thin-layer NGM plate under EVOS M5000. In the L1–L2 transition, animals were imaged continuously while GENTIS translocation was recorded. Bead-feeding assays were performed by exposing animals to fluorescent latex beads (Sigma, L3280) for 10 min and scoring intestinal bead uptake. Reversibility was tested by washing out FA after 10 min incubation and monitoring recovery of feeding. For the population-level pumping quiescence assay, animals that did not pump for 5 seconds were considered to be in pumping quiescence at the monitored time point. Similarly, animals that did not move for 5 seconds were defined as exhibiting locomotion quiescence at the monitored time point. Individual pumping frequency was measured over a 10 second interval.

## Statistics

Numerical data were analyzed using GraphPad Prism 10 Software (Graphpad, San Diego, CA) and presented as means ± S.E.M. unless otherwise specified, with *P* values calculated by unpaired two-tailed t-tests (comparisons between two groups), one-way ANOVA (comparisons across more than two groups) and two-way ANOVA (interaction between genotype and treatment), with post-hoc Tukey and Bonferroni’s corrections.

## Structural prediction and analysis

Structural modeling was performed using the Chai Discovery platform(*52*) under two conditions. In the first condition, the VHA-13 monomer was modeled in complex with one ATP molecule and six FA molecules. In the second condition, a VHA-12/VHA-13 heterodimer was modeled with one ATP and six FA molecules. For each condition, five independent prediction results were generated. The predicted complexes were examined in UCSF ChimeraX(*53*) to analyze the spatial relationship between FA and the ATP-binding pocket, and the electrostatic environment surrounding ATP and FA. The predicted ATP-binding site was consistent with that observed in homologous structures, and one potential FA-binding pocket was identified adjacent to the ATP-binding region.

## Acknowledgment

Some strains were provided by the *Caenorhabditis* Genetics Center (CGC), which is funded by the NIH Office of Research Infrastructure Programs (P40 OD010440), and by Drs. Henrik Bringmann, Josh Kaplan, and Matt Nelson. We also thank the *C. elegans* Reverse Genetics Core Facility (University of British Columbia), National Bioresource Project (Tokyo Women’s Medical University), Wormbase.org (NIH grant #U24 HG002223 to P. Sternberg), Wormatlas.org (NIH grant #OD010943 to D.H. Hall.), wormseq.org (Dr. E. O’Rourke), Aging Atlas (Dr. M. Wang) and CenGen for invaluable resources. The work was supported by NIH grants (R35GM139618 to D.K.M.), AHA (24TPA1288391) and 2025 UCSF PBBR New Frontier Research Award (D.K.M.)

## Author Contributions

Z.J. designed, performed and analyzed most experiments in this work, including characterization of GENTIS properties, live reporter imaging and phenotypic assays. J.L., B.W., Y.B. and S.W. helped on GENTIS transgenic construction, v-ATPase biochemical and imaging analyses. X.S., W.Z., C.C., Z.M. contributed to the design and plasmid construction of GENTIS. J.Z. modelled the structure of VHA-13 and its binding to FA. D.K.M. designed and analyzed the *C. elegans* experiments, contributed to project conceptualization, funding acquisition and editing the manuscript. All authors contributed to research materials, project conceptualization and editing manuscript.

---

**Fig. S1.**
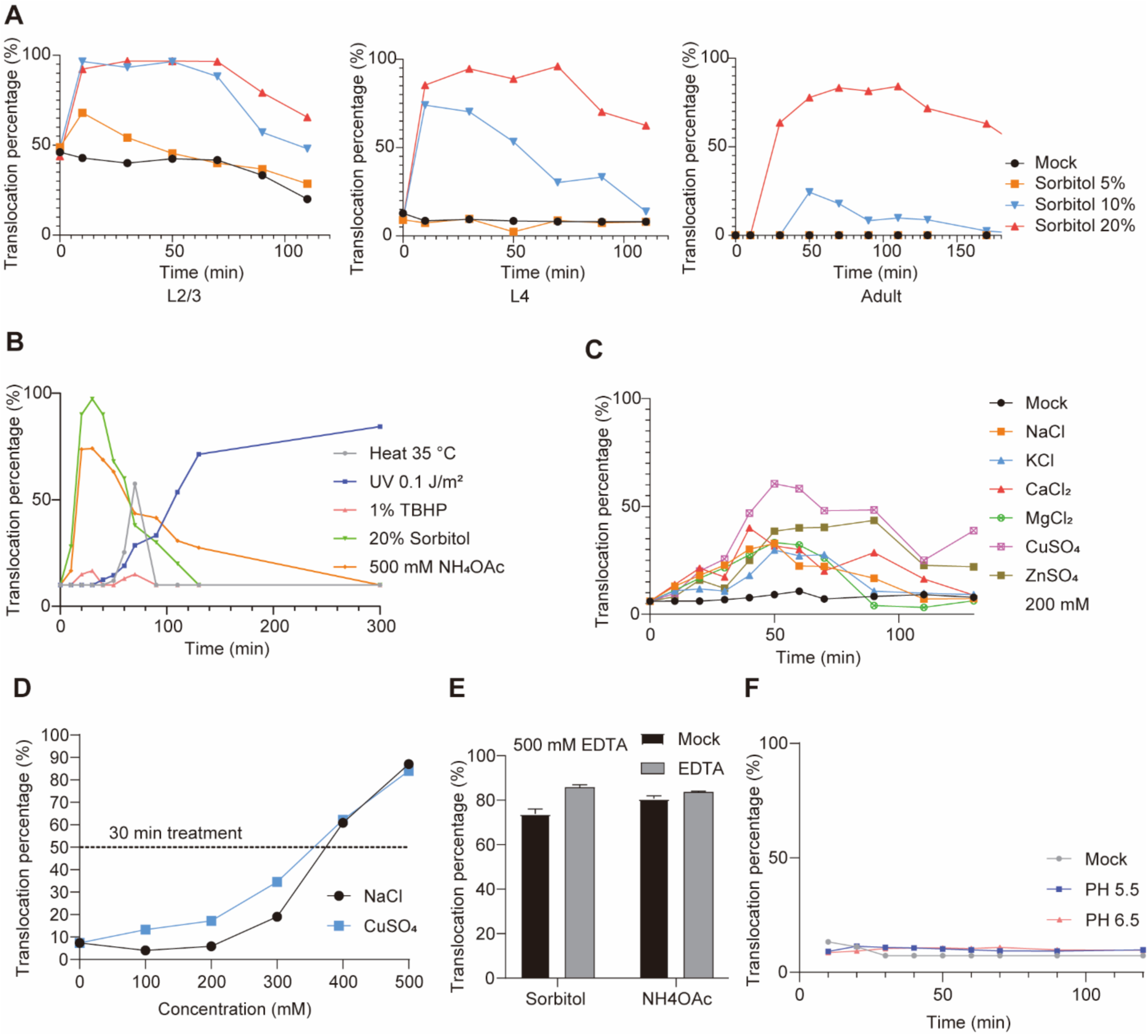
GENTIS sensor response kinetics and specificity under stress. (A) Time course of GENTIS nuclear translocation in populations of L2/L3, L4, and adult worms exposed to increasing concentrations of sorbitol (5%, 10%, and 20%). The percentage of animals showing nuclear localization was quantified over time. (B) GENTIS translocation under various stress conditions, including heat (35 °C), UV irradiation (0.1 J m⁻²), oxidative stress (1% TBHP), 20% sorbitol, and 500 mM NH_4_OAc. (C) Translocation percentages after exposure to different 200 mM metal ion solutions (NaCl, KCl, CaCl_2_, MgCl_2_, CuSO_4_, ZnSO_4_). (D) Dose-dependent response to NaCl and CuSO4 after 30 min incubation. (E) GENTIS nuclear import induced by 20% sorbitol or 500 mM NH_4_OAc was not affected by the metal chelator EDTA (500 mM). (F) Acidic environments (pH 5.5 and 6.5) did not induce measurable GENTIS translocation. For all assays, > 30 worms were quantified per condition. Error bars represent S.E.M.

**Fig. S2.**
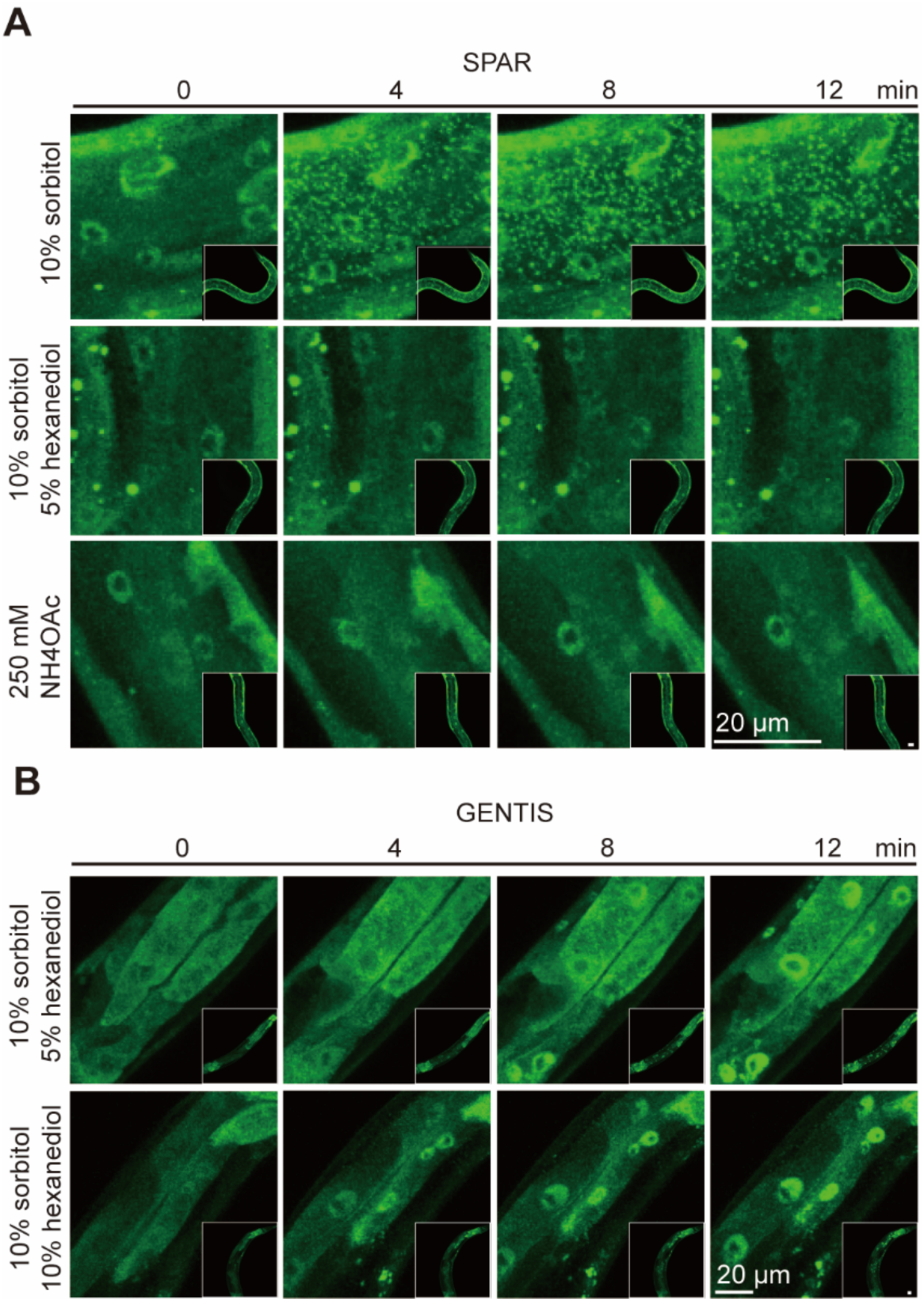
GENTIS translocation is independent of phase separation. (A) Representative high-resolution confocal images of worms expressing the SPAR reporter under hypertonic or high ionic strength conditions, with or without a phase-separation inhibitor. (B) Representative high-resolution images of worms expressing GENTIS under a hypertonic condition in the presence of the phase-separation inhibitor. Scale bars, 20 μm.

**Fig. S3.**
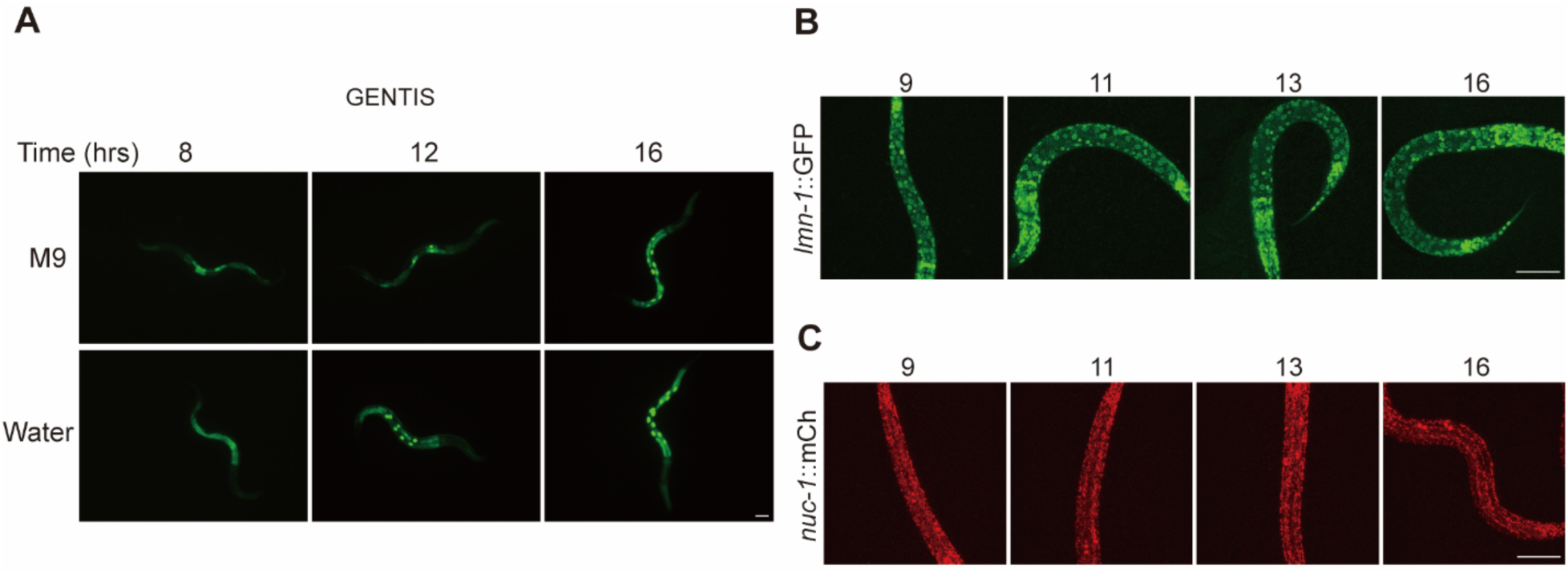
GENTIS translocation is independent of the changes in environmental tonicity and subcellular organelle markers during the L1–L2 molt. (A) GENTIS nuclear translocation during the L1–L2 molting stage is not inhibited by hypotonic pure water treatment. (B) Representative images of the nuclear envelope marker LMN-1::GFP at different time points during the L1–L2 molt. (C) Representative images of the lysosomal marker NUC-1::mCherry during the L1–L2 molt. Scale bars, 20 μm.

**Fig. S4.**
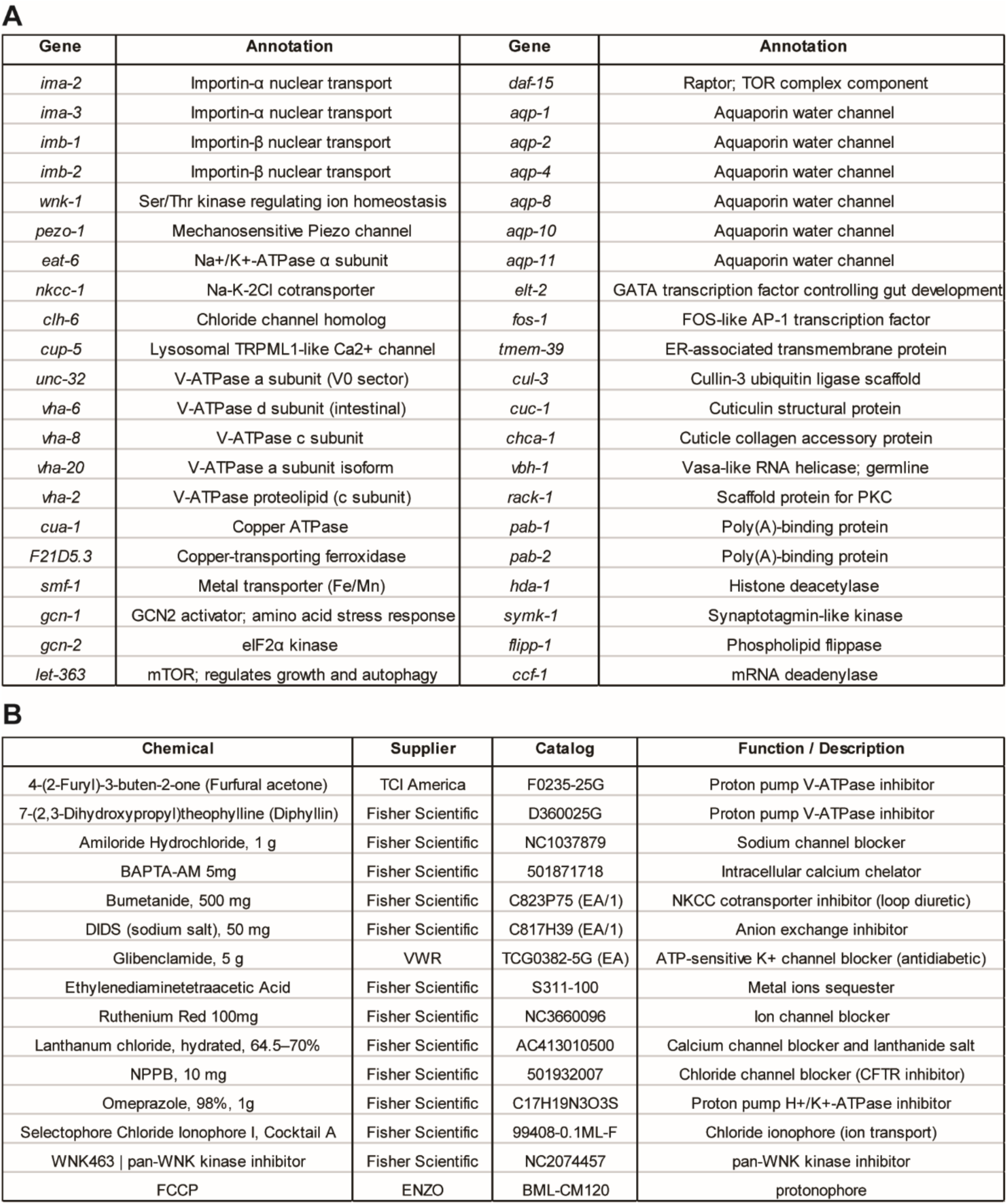
RNAi and pharmacological screening for regulators of GENTIS translocation. (A) List of genes targeted by RNAi for functional screening of GENTIS translocation under ionic stress. (B) List of small-molecule inhibitors and modulators tested for their effects on GENTIS translocation. The table summarizes compound name, supplier, catalog number, and known target or function.

**Fig. S5.**
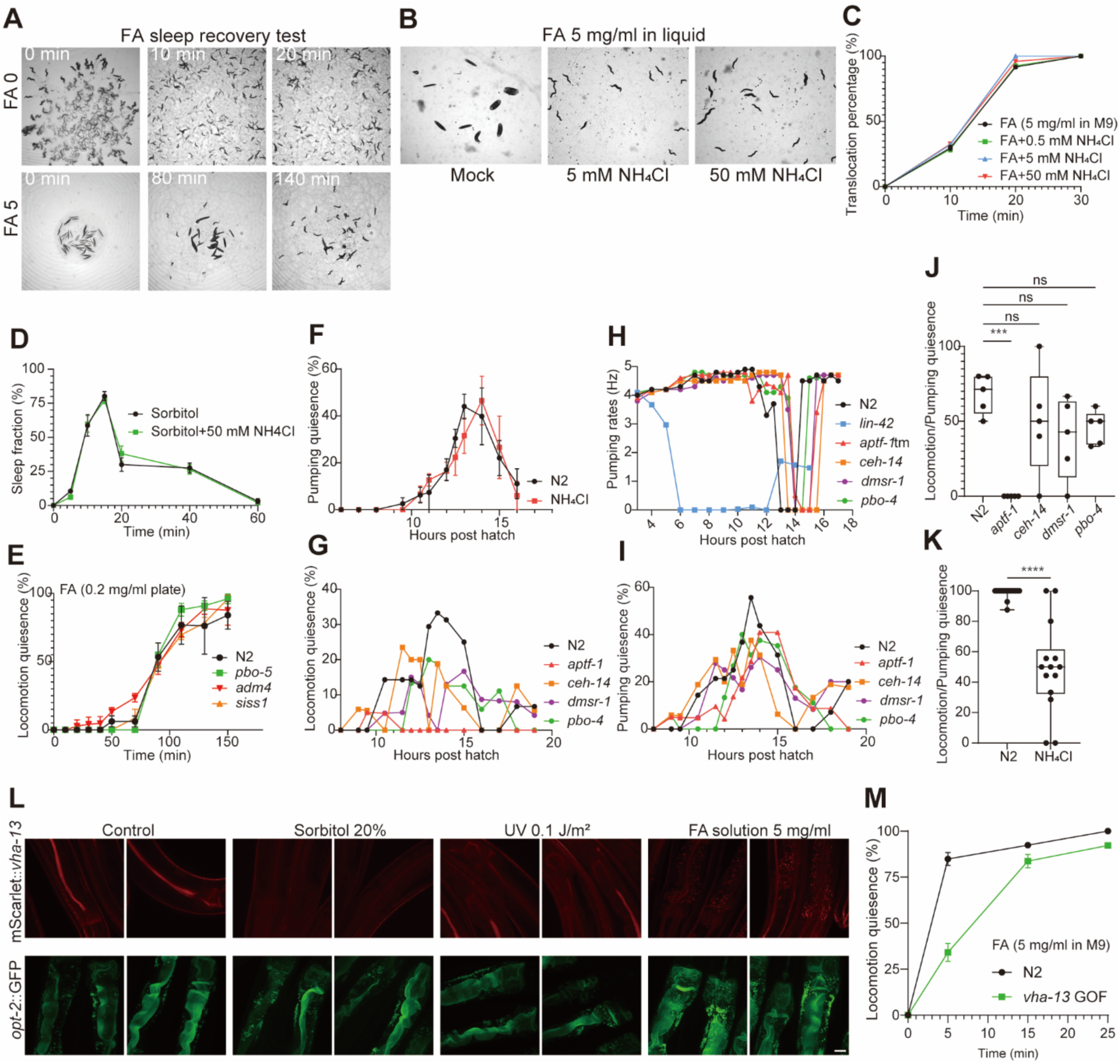
Reversibility, buffering, and genetic modulation of sleep and VHA-13 dynamics in *C. elegans*. (A) Representative images from the FA sleep recovery assay acquired using the WormLab imaging system. Animals treated with FA (5 mg/ml) recovered by 140 min after removal of FA. Control animals (0 mg/ml) remained active throughout. (B) Representative images of worms treated with FA (5 mg/ml) for 10 min in the presence of varying concentrations of NH_4_Cl, acquired using the WormLab imaging system. (C) GENTIS translocation is not affected by NH_4_Cl co-treatment, consistent with GENTIS responding to overall ionic strength rather than specifically to cytosolic proton stress. (D) Sleep fraction in worms treated with 20% sorbitol alone or in combination with 50 mM NH_4_Cl. NH_4_Cl co-treatment does not suppress sorbitol-induced sleep. (E) Sleep fraction (locomotion) over time in *pbo-5*, *adm-4*, and *siss-1* mutants following FA treatment (5 mg/ml in liquid). Sleep dynamics were comparable to wild-type (N2), indicating these genes are dispensable for FA-induced sleep. (F) Fraction of animals exhibiting pumping quiescence during the L1–L2 transition under NH_4_Cl or mock treatment. More than 20 worms were scored per plate, with three independent plates per condition. (G) Fraction of animals exhibiting locomotion quiescence during the L1–L2 transition in wild-type and mutant strains. (H) Individual pharyngeal pumping traces during L1–L2 transition for wild-type and sleep-related mutants. Indicated mutants displayed delayed sleep onset; *lin-42* mutants exhibited prolonged quiescence. (I) Fraction of animals exhibiting pumping quiescence during the L1–L2 transition in wild-type and mutant strains. (J) Ratio of locomotion to pumping quiescence during the L1–L2 molting sleep in wild-type and sleep-defective mutants. (K) Ratio of locomotion to pharyngeal pumping quiescence in wild-type animals with or without NH_4_Cl treatment during the L1–L2 transition. Each dot represents a timepoint during molting sleep. Three worms were analyzed per condition (J and K). (L) Apical mScarlet::VHA-13 signal and membrane-localized OPT-2::GFP in the anterior intestine under control conditions or following stress (20% sorbitol, 0.1 J/m^2^ UV, or 5 mg/ml FA). VHA-13 signal is reduced in all stress conditions, whereas OPT-2::GFP remains intact, indicating that membrane structure is preserved while proton pump levels decline. (M) FA-induced sleep is attenuated in *vha-13* gain-of-function (GOF) mutants. Locomotion quiescence in response to 5 mg/ml FA (in M9 buffer) was delayed in *vha-13* GOF animals compared to wild type (N2). More than 30 worms were quantified per condition, and experiments were repeated independently three times. Data represent mean ± S.E.M.

## References

1. S. S. Campbell, I. Tobler, Animal sleep: a review of sleep duration across phylogeny. Neurosci Biobehav Rev 8, 269–300 (1984).

2. K. Singh, J. Y. Ju, M. B. Walsh, M. A. DiIorio, A. C. Hart, Deep conservation of genes required for both Drosphila melanogaster and Caenorhabditis elegans sleep includes a role for dopaminergic signaling. Sleep 37, 1439–1451 (2014).

3. H. Huang, Y. Zhu, M. N. Eliot, V. S. Knopik, J. E. McGeary, M. A. Carskadon, A. C. Hart, Combining Human Epigenetics and Sleep Studies in Caenorhabditis elegans: A Cross-Species Approach for Finding Conserved Genes Regulating Sleep. Sleep 40, zsx063 (2017).

4. R. C. Anafi, M. S. Kayser, D. M. Raizen, Exploring phylogeny to find the function of sleep. Nat Rev Neurosci 20, 109–116 (2019).

5. J. M. Siegel, Sleep function: an evolutionary perspective. Lancet Neurol 21, 937–946 (2022).

6. R. Lakhiani, S. Shanavas, K. Melnattur, Comparative biology of sleep in diverse animals. J Exp Biol 226, jeb245677 (2023).

7. T. Worth, Sleep is essential - researchers are trying to work out why. Nature, doi: 10.1038/d41586-025-00964-w (2025).

8. N. F. Trojanowski, D. M. Raizen, Call it Worm Sleep. Trends Neurosci 39, 54–62 (2016).

9. R. C. Cassada, R. L. Russell, The dauerlarva, a post-embryonic developmental variant of the nematode Caenorhabditis elegans. Dev Biol 46, 326–342 (1975).

10. D. M. Raizen, J. E. Zimmerman, M. H. Maycock, U. D. Ta, Y. You, M. V. Sundaram, A. I. Pack, Lethargus is a Caenorhabditis elegans sleep-like state. Nature 451, 569–572 (2008).

11. J. Schwarz, I. Lewandrowski, H. Bringmann, Reduced activity of a sensory neuron during a sleep-like state in Caenorhabditis elegans. Curr Biol 21, R983–984 (2011).

12. S. Iwanir, N. Tramm, S. Nagy, C. Wright, D. Ish, D. Biron, The microarchitecture of C. elegans behavior during lethargus: homeostatic bout dynamics, a typical body posture, and regulation by a central neuron. Sleep 36, 385–395 (2013).

13. J. Y. Cho, P. W. Sternberg, Multilevel modulation of a sensory motor circuit during C. elegans sleep and arousal. Cell 156, 249–260 (2014).

14. M. Turek, I. Lewandrowski, H. Bringmann, An AP2 transcription factor is required for a sleep-active neuron to induce sleep-like quiescence in C. elegans. Curr Biol 23, 2215–2223 (2013).

15. R. D. Nath, E. S. Chow, H. Wang, E. M. Schwarz, P. W. Sternberg, C. elegans Stress-Induced Sleep Emerges from the Collective Action of Multiple Neuropeptides. Curr Biol 26, 2446–2455 (2016).

16. A. J. Hill, R. Mansfield, J. M. N. G. Lopez, D. M. Raizen, C. Van Buskirk, Cellular stress induces a protective sleep-like state in C. elegans. Curr Biol 24, 2399–2405 (2014).

17. M. D. Nelson, K. H. Lee, M. A. Churgin, A. J. Hill, C. Van Buskirk, C. Fang-Yen, D. M. Raizen, FMRFamide-like FLP-13 neuropeptides promote quiescence following heat stress in Caenorhabditis elegans. Curr Biol 24, 2406–2410 (2014).

18. J. Konietzka, M. Fritz, S. Spiri, R. McWhirter, A. Leha, S. Palumbos, W. S. Costa, A. Oranth, A. Gottschalk, D. M. Miller, A. Hajnal, H. Bringmann, Epidermal Growth Factor Signaling Promotes Sleep through a Combined Series and Parallel Neural Circuit. Curr Biol 30, 1–16.e13 (2020).

19. L. Rossi, K. Amoako, I. Busack, L. Golinelli, A. Courtney, J. Besseling, W. Schafer, I. Beets, H. Bringmann, The neuropeptide FLP-11 induces and self-inhibits sleep through the receptor DMSR-1 in Caenorhabiditis elegans. Curr Biol 35, 2183–2194.e10 (2025).

20. M. J. Iannacone, I. Beets, L. E. Lopes, M. A. Churgin, C. Fang-Yen, M. D. Nelson, L. Schoofs, D. M. Raizen, The RFamide receptor DMSR-1 regulates stress-induced sleep in C. elegans. Elife 6, e19837 (2017).

21. A. J. Hill, B. Robinson, J. G. Jones, P. W. Sternberg, C. Van Buskirk, Sleep drive is coupled to tissue damage via shedding of Caenorhabditis elegans EGFR ligand SISS-1. Nat Commun 15, 10886 (2024).

22. C. Van Buskirk, P. W. Sternberg, Epidermal growth factor signaling induces behavioral quiescence in Caenorhabditis elegans. Nat Neurosci 10, 1300–1307 (2007).

23. C. Cirelli, Sleep, synaptic homeostasis and neuronal firing rates. Curr Opin Neurobiol 44, 72–79 (2017).

24. T. Porkka-Heiskanen, R. E. Strecker, M. Thakkar, A. A. Bjorkum, R. W. Greene, R. W. McCarley, Adenosine: a mediator of the sleep-inducing effects of prolonged wakefulness. Science 276, 1265–1268 (1997).

25. R. Sarnataro, C. D. Velasco, N. Monaco, A. Kempf, G. Miesenböck, Mitochondrial origins of the pressure to sleep. Nature 645, 722–728 (2025).

26. S. Hajjar, X. Zhou, pH sensing at the intersection of tissue homeostasis and inflammation. Trends Immunol 44, 807–825 (2023).

27. P. Mitchell, J. Moyle, Chemiosmotic Hypothesis of Oxidative Phosphorylation. Nature 213, 137–139 (1967).

28. M. Forgac, Vacuolar ATPases: rotary proton pumps in physiology and pathophysiology. Nat Rev Mol Cell Biol 8, 917–929 (2007).

29. R. Sarnataro, C. D. Velasco, N. Monaco, A. Kempf, G. Miesenböck, Mitochondrial origins of the pressure to sleep. Nature 645, 722–728 (2025).

30. W.-Z. Zeng, T.-L. Xu, Proton production, regulation and pathophysiological roles in the mammalian brain. Neurosci Bull 28, 1–13 (2012).

31. A. A. Beg, G. G. Ernstrom, P. Nix, M. W. Davis, E. M. Jorgensen, Protons Act as a Transmitter for Muscle Contraction in *C. elegans*. Cell 132, 149–160 (2008).

32. S.-A. Li, X.-Y. Meng, Y.-J. Zhang, C.-L. Chen, Y.-X. Jiao, Y.-Q. Zhu, P.-P. Liu, W. Sun, Progress in pH-Sensitive sensors: essential tools for organelle pH detection, spotlighting mitochondrion and diverse applications. Front Pharmacol 14, 1339518 (2023).

33. A. M. M. Gest, A. Z. Sahan, Y. Zhong, W. Lin, S. Mehta, J. Zhang, Molecular Spies in Action: Genetically Encoded Fluorescent Biosensors Light up Cellular Signals. Chem Rev 124, 12573–12660 (2024).

34. Q.-Y. Zhang, S. Mehta, J. Zhang, Recent advances in spying on cell signaling with fluorescent biosensors. Curr Opin Cell Biol 95, 102546 (2025).

35. G. De Crescenzo, J. R. Litowski, R. S. Hodges, M. D. O’Connor-McCourt, Real-Time Monitoring of the Interactions of Two-Stranded de Novo Designed Coiled-Coils: Effect of Chain Length on the Kinetic and Thermodynamic Constants of Binding. Biochemistry 42, 1754–1763 (2003).

36. A. P. Jalihal, S. Pitchiaya, L. Xiao, P. Bawa, X. Jiang, K. Bedi, A. Parolia, M. Cieslik, M. Ljungman, A. M. Chinnaiyan, N. G. Walter, Multivalent Proteins Rapidly and Reversibly Phase-Separate upon Osmotic Cell Volume Change. Mol Cell 79, 978–990.e5 (2020).

37. X. Shu, Imaging dynamic cell signaling *in vivo* with new classes of fluorescent reporters. Current Opinion in Chemical Biology 54, 1–9 (2020).

38. Q. Zhang, H. Huang, L. Zhang, R. Wu, C.-I. Chung, S.-Q. Zhang, J. Torra, A. Schepis, S. R. Coughlin, T. B. Kornberg, X. Shu, Visualizing Dynamics of Cell Signaling In Vivo with a Phase Separation-Based Kinase Reporter. Molecular Cell 69, 334–346.e4 (2018).

39. W. Cheng, W. Dai, W. Chen, H. Xue, Z. Zhao, Z. Jiang, H. Li, J. Liu, F. Huang, M. Cai, L. Zheng, Z. Yu, D. Peng, J. Zhang, Nematodes exposed to furfural acetone exhibit a species-specific vacuolar H+-ATPase response. Ecotoxicology and Environmental Safety 288, 117407 (2024).

40. J. Goudeau, C. S. Sharp, J. Paw, L. Savy, M. D. Leonetti, A. G. York, D. L. Updike, C. Kenyon, M. Ingaramo, Split-wrmScarlet and split-sfGFP: tools for faster, easier fluorescent labeling of endogenous proteins in Caenorhabditis elegans. Genetics 217, iyab014 (2021).

41. A. Kempf, S. M. Song, C. B. Talbot, G. Miesenböck, A potassium channel β-subunit couples mitochondrial electron transport to sleep. Nature 568, 230–234 (2019).

42. A. Vaccaro, Y. Kaplan Dor, K. Nambara, E. A. Pollina, C. Lin, M. E. Greenberg, D. Rogulja, Sleep Loss Can Cause Death through Accumulation of Reactive Oxygen Species in the Gut. Cell 181, 1307–1328.e15 (2020).

43. R. Zhou, K. Li, X. Hu, S. Fan, Y. Gao, X. Xue, Y. Bu, H. Zhang, Y. Wang, C. Wei, S. Zhang, Z. Xie, C. Liu, P. Chen, Z. Yin, D. Ren, Sleep Deprivation Activates a Conserved Lactate-H3K18la-RORα Axis Driving Neutrophilic Inflammation Across Species. Adv Sci (Weinh) 12, e04028 (2025).

44. D. Sang, K. Lin, Y. Yang, G. Ran, B. Li, C. Chen, Q. Li, Y. Ma, L. Lu, X.-Y. Cui, Z. Liu, S.-Q. Lv, M. Luo, Q. Liu, Y. Li, E. E. Zhang, Prolonged sleep deprivation induces a cytokine-storm-like syndrome in mammals. Cell 186, 5500–5516.e21 (2023).

45. E. Allman, D. Johnson, K. Nehrke, Loss of the apical V-ATPase a-subunit VHA-6 prevents acidification of the intestinal lumen during a rhythmic behavior in C. elegans. Am J Physiol Cell Physiol 297, C1071–1081 (2009).

46. I. Dimov, M. F. Maduro, The C. elegans intestine: organogenesis, digestion, and physiology. Cell Tissue Res 377, 383–396 (2019).

47. S. Iwanir, N. Tramm, S. Nagy, C. Wright, D. Ish, D. Biron, The microarchitecture of C. elegans behavior during lethargus: homeostatic bout dynamics, a typical body posture, and regulation by a central neuron. Sleep 36, 385–395 (2013).

48. P. M. Kane, Disassembly and Reassembly of the Yeast Vacuolar H+-ATPase in Vivo(*). Journal of Biological Chemistry 270, 17025–17032 (1995).

49. T. Vasanthakumar, K. A. Keon, S. A. Bueler, M. C. Jaskolka, J. L. Rubinstein, Coordinated conformational changes in the V1 complex during V-ATPase reversible dissociation. Nat Struct Mol Biol 29, 430–439 (2022).

50. J. F. Staples, Metabolic suppression in mammalian hibernation: the role of mitochondria. J Exp Biol 217, 2032–2036 (2014).

51. S. Brenner, The genetics of Caenorhabditis elegans. Genetics 77, 71–94 (1974).

52. C. Discovery, J. Boitreaud, J. Dent, M. McPartlon, J. Meier, V. Reis, A. Rogozhnikov, K. Wu, Chai-1: Decoding the molecular interactions of life. bioRxiv [Preprint] (2024). 10.1101/2024.10.10.615955.

53. E. F. Pettersen, T. D. Goddard, C. C. Huang, E. C. Meng, G. S. Couch, T. I. Croll, J. H. Morris, T. E. Ferrin, UCSF ChimeraX: Structure visualization for researchers, educators, and developers. Protein Science 30, 70–82 (2021).

